# Using Imaris to rigorously track PET-defined sites of lung inflammation in *Mycobacterium tuberculosis*-exposed non-human primates

**DOI:** 10.1101/2025.07.04.663191

**Authors:** Estefania Hurtado, Xavier Alvarez, Deepak Kaushal, Smriti Mehra, Vitaly V. Ganusov

## Abstract

Aerosol exposure of non-human primates (**NHPs**) to *Mycobacterium tuberculosis* (**Mtb**) typically results in discrete sites of inflammation of the lung that is detectable by 2-deoxy-2-[fluorine-18]fluoro-D-glucose (^18^F-**FDG**)-based PET/CT scans. Such scans are often analyzed using software such as Invicro VivoQuant or OsiriX as 3D images by manual labeling sites of PET signal using 2D slices and by reporting maximal standardized uptake value (**SUV**_max_) either of the whole lung or of individual lesions. Here we propose a pipeline for analysis of the same PET/CT scans using Imaris, a proprietary software typically used for analysis of fluorescent microscopy data. We show that by using locations of spine vertebra (denoted as “landmarks”) we can align serials scans of the same animal, and by using automated (with some manual corrections) image segmentation of PET scans in 3D as “surfaces”, we can accurately define location of all sites of inflammation in the lung and lung-associated thoracic lymph nodes (**LNs**). We show that there is an excellent correlation between individual lesion’s SUV_max_ determined by VivoQuant and maximum intensity determined by Imaris suggesting utility of this approach. Imaris also provides wealth of additional information for each of the identified lesions such as volume, location, shape, surface area, and others, and each lesion can be exported in Virtual Reality file format (.wrl) allowing for detailed and rigorous analyses of how features of these PET-defined lesions evolve over time and correlate with the outcome of infection and/or treatment.

## Introduction

Tuberculosis (**TB**), a disease caused by *Mycobacterium tuberculosis* (**Mtb**) bacilli, is the leading cause of death from a single infectious as of 2023 (ref ^1^). There were over 10 million cases and over 1.2 million deaths due to TB in 2023, and a quarter of the world’s population has evidence of past or present Mtb infection^1,2^. Despite high numbers of TB cases and TB-related deaths, humans are relatively resistant to TB as some estimates suggest that only 5-10% of Mtb-exposed individuals progress to active disease^3–6^; cohort studies suggested more variable proportions, e.g., from *<* 2% in pre-antibiotic era in the UK to 14.5% more recently in Australia^7,8^ Factors, determining why upon exposure to Mtb some individuals progress to TB while others are able to control (and perhaps clear) the infection remain poorly understood^9,10^; yet, children and immunocompromised individuals (e.g., people living with HIV) have a higher risk of progressing to TB upon Mtb exposure^8,11^.

Imaging, specifically X-rays and computed tomography (**CT**), has been a cornerstone of TB control programs as detecting abnormalities in the lung or other tissues is often required (along with Mtb positive culture from tissue samples) to diagnose TB^12,13^. Such imaging technology has been key for understanding how Mtb-infected individuals progress to TB in cohort studies ^7^ and has helped detect a high proportion of individuals with signs of lung disease and yet without obvious TB-associated symptoms^14–18^. However, detecting early signs of progression to TB with CT may be difficult as changes in the tissue structure, detectable by X-rays, may take time to develop ^12^. Therefore, other lung imaging techniques such as positron emission tomography (**PET**) have been introduced to gain better insights into TB pathogenesis.

The key element of PET is the injection of radiotracer that has some specificity for the target of interest and is modified to emit positrons allowing to detect the 3D locations of the radio-tracer molecules. Perhaps the most commonly used radiotracer is the 2-deoxy-2-[fluorine-18]fluoro-D-glucose (^18^F-FDG or simply **FDG**) that is glucose analog and is taken up by body cells with high metabolic activity^19^. In PET/CT scanners, CT then allows to image the overall body composition while FDG-PET allows to image sites of high cellular metabolic activity^20^. The fact that FDG-PET allows to detect, for example, rapidly dividing cells has been extensively used in detecting solid cancers and track response of cancers to treatment^21,22^.

FDG-PET/CT has been also used to understand TB pathogenesis. In humans, to speed up development of novel treatments, PET/CT has been used to detect impact of various drug combinations on lung inflammation in treated TB patients^23–26^. In animals, PET/CT has been also used to understand response of Mtb infection to drug treatment, to study disease progression in Mtb-infected animals, including following co-infection with SIV and during antiretroviral treatment (**ART**), and to evaluate impact of vaccination on protection against TB development^27–37^. PET radiotracers that are more specific for Mtb have been also developed^38^. Yet, because of high costs of equipment and consumables, and of expertise involved in performing PET/CT scans (e.g., a need to rapidly generate radioactive tracer), the use of PET/CT in TB has been primarily at pre-clinical stages^26^.

PET/CT scans of monkeys generate data in Digital Imaging and Communications in Medicine (**DICOM**) format that is then read by specialized DICOM file viewers such as Invicro VivoQuant or OsiriX that typically are bundled with the PET/CT scanner^39,40^. The resulting PET scans are then typically analyzed by manually labeling areas of high PET signal, indicating sites of inflammation; CT signal is then used to confirm location of individual lesions. For each identified site of inflammation (and for each scan) it is typical to report maximal standardized uptake value (**SUV**_max_) indicating the relative degree of radiotracer uptake. Change in lung SUV_max_ with time has been used to indicate control of Mtb infection, e.g., in vaccinated monkeys or monkeys co-infected with SIV^30,31,41–43^.

Because there may be tens of sites of inflammation in lungs of Mtb-infected monkeys, manual labeling of each site of inflammation in PET/CT scans is laborious; in our experience it may take one day to carefully label one scan. Results may also depend on how the operator defines boundaries for each identified site/lesion. Furthermore, DICOM viewers typically only allow to analyze one scan at a time making it difficult to rigorously follow serial scans for the same subject due to minor shits occurring during body positioning on the scanner’s bench. DICOM viewers also deliver relatively limited additional information regarding the user-defined lesions, e.g., typically only SUV_max_ and volume per lesion is provided. Here we introduce a novel pipeline to analyze PET/CT scans of macaque lungs in Imaris, a proprietary software commonly used to analyze microscopy data. Upon import of DICOM scans (CT + PET) in Imaris, scan signals are converted into channel intensities. By using locations of three elements for each of two thoracic vertebra (T2 and T10), called landmarks, we align scans of the same animal done at different time points, thus allowing for more rigorous comparison of locations different sites of inflammation. By using the Surfaces tool in Imaris, we then use a single threshold value for PET signal intensity (sometimes the threshold may vary between different scans) to identify all key inflammation sites/lesions and use 3D Imaris viewer to carefully manually edit identified Surfaces. Importantly, Imaris provides a wealth of information for each of the defined site of inflammation such as their 3D coordinates (including relative to landmarks), volume, surface area, sphericity, and intensity of both channels; by using 12 scans from 4 animals we show that there is an excellent correspondence between maximum channel intensity *I*_max_, provided by Imaris, and SUV_max_ provided by DICOM viewer VivoQuant (with some potential variability between individual scans). Visualization of lesion location and their properties (e.g., volume) allow to track development of TB over time, thus, paving the way for more rigorous understanding why some Mtb-infected monkeys progress to active disease while others control the infection.

## Materials & Methods

### Data

We analyzed PET/CT imaging data from two different studies: animals 41634 and 40884 were from our previous study^34^, while animals 41883 and 44104 were from a study that has not yet been published (Singh et al. (in prep)). The analysis was for four animals, two males and two females, animal IDs are 41883 (M), 44104 (F), 41634 (M), and 40884 (F). Scans were performed at 11, 16, and 22 weeks post Mtb infection for 41883 and 44104, and at 6, 14, and 18 weeks post-infection in 41634 and 40884. Other details are as follows

- ID 41883. Infection dose: 10 CFU of CDC1551. Animal was co-infected with SIV (at 9 wks post Mtb infection). Animal had signs of active disease starting at wk 11.
- ID 44104. Infection dose: 10 CFU of CDC1551. Animal was co-infected with SIV (at 9 wks post Mtb infection). Animal had no disease at wk 11 but active disease at wk 16 and 22.
- ID 41634. Infection dose: 25-50 CFU. Animal had no disease at wk 6 or 14 but active disease at wk 18.
- ID 40884. Infection dose: 25-50 CFU. The animal did not have disease at all scan time points.

Disease status was defined based on the collection of markers including symptoms and blood-based indicators (e.g., rapid rise in C-reactive protein concentration^44^).

### Imaging procedures

PET/CT imaging of non-human primates at Texas Biomed follows standard operating procedures (**SOP**) protocol 425.01 “RAM Handling and Animal Procedures in PET-CT Imaging”. We use Mediso LFER150 PET/CT scanner (Budapest)^45^. For CT imaging we use a breath-hold protocol as per the instructions in the PET-CT manual of the scanner. We then check the quality of the acquired scans for the inclusion of the region of interest, diaphragmatic breathing movements, animal position/tilt, and resolution. If needed and after consultation and approval of the veterinary team, we perform one more breath-hold CT scan. After the CT scan is completed, we ensure that the animal is comfortable before starting the PET acquisition. We use the standard radio tracer dose, 1 mCi/kg of 18F-FDG, that is calculated using the weight of the study animal. The dose is delivered in 3ml in a 5ml syringe. After injection of radioisotope, the animal is rested and observed clinically for the next 20-30 minutes, to allow for the uptake of 18F-FDG glucose. We perform a static PET scan as per the instructions in the PET-CT manual about 45 minutes after 18F-FDG injection. For scan reconstruction we used The 3D Tera-Tomo engine reconstruction method provided by the Meduso (the files are processed and uploaded to our specialized Meduso server). Parameters of reconstruction were those recommended for non-human primates by Mediso, specifically voxel large 426×426×425 *µ*m. PET emission data were reconstructed with isotropic voxels of 0.8×0.8×0.8 mm to balance resolution and signal-to-noise ratios ^45^.

### VivoQuant-based lesion quantification

We used PET/CT DICOM file viewer Invicro VivoQuant (https://vivoquant.com/) for quan-tification of SUV from each of the lesions. We identified lesions in the scans (CT and/or PET) by going through each of Z-stacks and using XY-plane slicer in the whole lung. We then highlighted the lesion area for each of the 2D slice through each lesion. Then for each lesion we calculated specific label uptake, characterized by SUV, and export these values into.csv file format.

### Imaris-based lesion quantification

Imaris is a software used to rigorously analyze data from microscopy experiments, typically from confocal or two-photon microscopes. However, more recently Imaris is able to import other types of imaging data, including from DICOM files from PET/CT scans, by creating channels for each of the modalities in the scan (i.e., separate channels for CT and PET). In this way, Imaris assigns values for channel intensity that may not fully match the PET values interpreted by the standard DICOM viewers and thus may need to be adjusted using information on the injected dose of the label, weight of the animal, etc. Imaris also allows to import serial scans of the same animal into the same file as different time frames thus allowing to more accurately connect lesions identified at different times in the same animal. We now use a feature in Imaris of drift correction that allows to rigorously align scans of the same animal done at different times. We use landmarks – locations of six different locations of vertebra found in CT channel to align the scans. Imaris also has an automated function of creating a 3D object “Surface” around a specific region of interest. In short, one would define a threshold intensity which would define the boundary of the lesion, and Imaris would then produce the 3D rendering of the lesion. The benefit of this process is that it is rapid and only depends on the threshold defined for the lesions in the scan (threshold may be also lesion-dependent), and thus is well-defined and thus can be reproduced by another user. Alternatively, one can identify multiple lesions by using one threshold value for the whole scan, and then one can edit the Surfaces objects to remove sites that are not expected to be associated with Mtb infection (e.g., spines). Imaris then provides information about every Surface produced including its location, volume, sphericity, total signal intensity, etc, and the Surface can be also exported in the standard Virtual Reality file format (.wrl) that can be analyzed further (e.g., to calculate distances between lesions, etc). We used the following formula to calculate the normalized PET channel lesion intensity *nI*_max_ given maximal raw intensity *I*_max_ provided by Imaris:

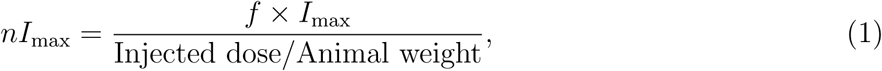

where *f* = 10^4^ is the scaling factor^46,47^. Because the values for the injected dose were relatively large in our data (∼ 10^8^), we used the scaling factor *f* to make values of *nI*_max_ comparable to typical SUV_max_ provided by the VivoQuant viewer. Ultimately, however, we are not concerned with the actual values of *nI*_max_ but rather how they vary between different lesions in a given scan or between different scans of the same animal; therefore, the actual value of the scaling parameter *f* does not influence our results. We are also exploring the possibility of exporting metadata from PET and CT DICOM files, specifically the intercepts and the slopes that convert voxel intensity into SUV or HU units, respectively.

## Results

### Analysis of PET/CT Images in Invicro VivoQuant and Imaris

Previous studies have outlined methodologies for analyzing PET/CT scans using DICOM viewers, such as Osirix or Invicro VivoQuant^39,40^; at Texas Biomed we use a somewhat similar pipeline with VivoQuant (**Figure 1**A). Our process consists of first acquiring the DICOM files for the PET/CT scans for each specific time point, and then converting the units from becquerels per milliliter to SUV (**Figure 1**Ai&ii). Once this conversion is finalized, a region of interest (**ROI**) is manually drawn for each lesion/site of inflammation (**Figure 1**Aiii). To complete this process, it is imperative to scroll slice by slice (z-direction) and across all three different views (coronal, sagittal, and transverse) in order to verify that the lesion is accurately tracked and its boundary lines are well defined. This is a long and time-consuming process that is highly dependent on the operator’s decision making when tracking these lesions (**Figure 2**Ai). Once all the ROIs are segmented, the.csv file is exported which contains basic metrics such as various measures of standardized uptake value (**SUV**) such as SUV_min_, SUV_max_, SUV_total_ as well as lesion volume and Hounsfield Units (**HUs**, **Figure 1**Aiv).

**Figure 1:**
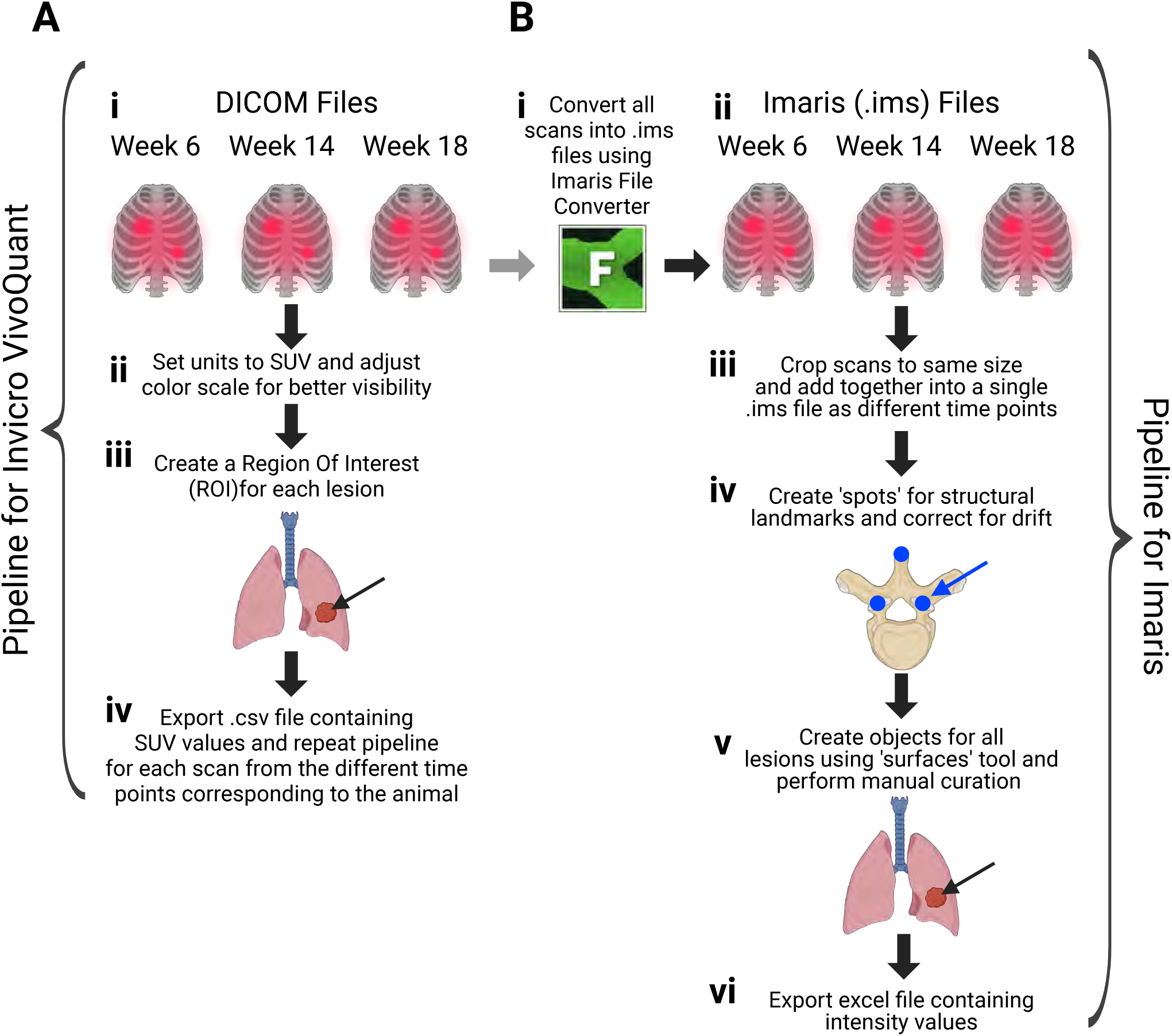
The pipeline of processing PET/CT imaging data with Invicro VivoQuant and Imaris. We illustrate how PET/CT scans are typically analyzed with Invicro VivoQuant, the standard proprietary packages for PET/CT data analysis (**A**), and with Imaris, proprietary packages typically used for fluorescent microscopy data analysis (**B**). **A**: With VivoQuant **(i-iv)**, the operator analyzes reconstructed 3D images by using 2D slices and manually labeling contours of the lesions, defined by PET signal. VivoQuant then provides various SUV characteristics, such as max SUV, for ether individual lesions or the whole lung. **B**: With Imaris **(i-vi)**, PET/CT DICOM files are imported as 2-channel images, and scans done at different times can be incorporated into the same Imaris file by adding new time frames. Then scans for different time points are aligned using “Correct for drift” routine by using landmarks (6 locations of spine vertebra). Then region of interest (**ROI**) is then identified around each suspected lesion and the data in ROI is processed using 3D routine “Surfaces”. All lesions then can be grouped into the specific group depending on the location of the lesion (e.g., lung vs. lymph nodes).

**Figure 2:**
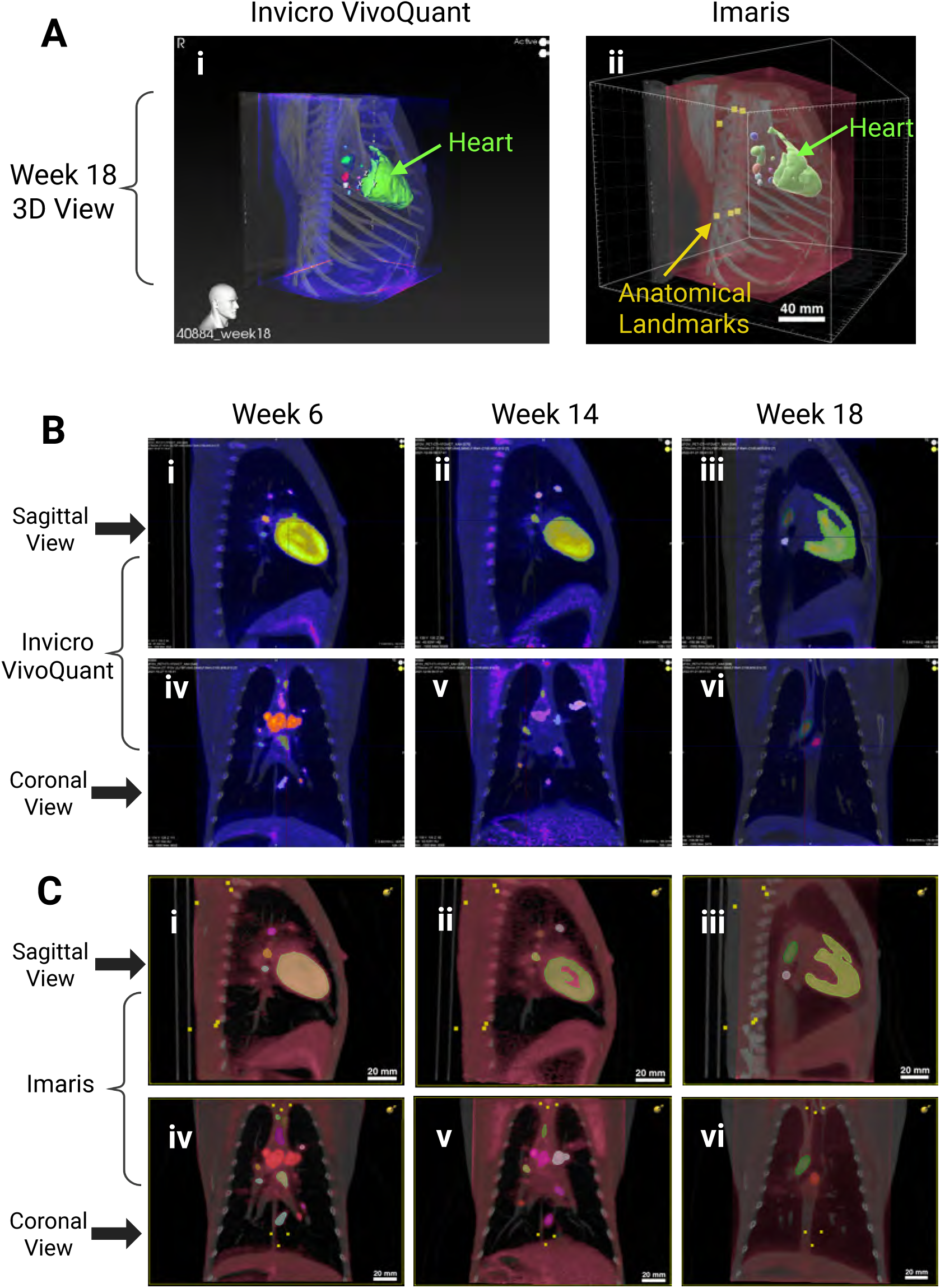
Lesions identified with the standard PET/CT software (Invicro VivoQuant) can be also traced using semi-automated tools in Imaris. Lesions identified for animal 40884 using VivoQuant (**Ai** and **B**) or Imaris (**Aii** and **C**) at different time points and different views. **A**: 3D view of lesions at 18 weeks post-infection in VivoQuant **(i)** and the same lesions in Imaris **(ii)**. **B-C**: Slice views of lesions identified with VivoQuant (**B**) or Imaris (**C**) at 6 **(i, iv)**, 14 **(ii, v)**, and 18 **(iii, vi)** weeks post-infection in both sagittal **(i-iii)** and coronal **(iv-vi)** slice views (see also **Supplemental Figures S1 and S2**). PET signal associated with the heart (shown by an arrow in **Ai-ii**) was excluded from our analyses.

At first glance, the analysis process with VivoQuant may seem relatively short, but it is actually quite time-consuming; depending on the complexity of the scan and expertise of operator in our experience it takes 4-8 hours to fully annotate one scan. In part, this is because creating ROIs in a DICOM viewer requires intensive manual scrolling through all 2D slices of the PET/CT scan to ensure that the entire area of inflammation is accurately defined with clear boundaries and margins. Also, because VivoQuant does not directly provide other information about identified lesions (e.g., their coordinates), we investigated whether using other software could help speeding up and more rigorously quantifying PET/CT scans.

Given our experience with analysis of imaging data from intravital microscopy experiments, we decided to use Imaris (https://imaris.oxinst.com/), a software package we have been using to track dynamics of Plasmodium sporozoites and liver-localized CD8 T cells in mice ^48–50^. We therefore developed a pipeline to track development of sites of inflammation from PET/CT scans in Mtb-infected non-human primates (**NHPs**, **Figure 1**B). In this pipeline, we use VivoQuant to export co-registered PET and CT scans separately and convert them into Imaris files (**Figure 1**Bi); these files are then imported as separate channels (CT – channel 1, PET – channel 2, **Figure 1**Bii). Importantly, Imaris allows to add images from different time points to the same file; we therefore, crop the converted images to the same size and import them as sequential time frames (**Figure 1**Biii). Image cropping is required to ensure that every scan across all time points has the same dimensions but it is important to note that only non-essential regions, such as background or artifacts, are cropped.

After merging all scans into a single file, we use the Imaris’ “Spots” tool to define the structural landmarks that should remain largely unchanged between the scans of the same animal (we selected those with the help of a veterinarian). For consistency, we selected two thoracic vertebrae separated by 6–8 segments, specifically T2 and T10. While other vertebrae could be used, the key criterion is a separation of 6–8 thoracic vertebrae between them. For each vertebra, three regions were marked as landmarks, totaling six anatomical reference points per scan. Two landmarks correspond to the tips of the spinous processes, while the remaining four represent the junctions of the pedicle, lamina, and transverse processes (**Figure 1**Biv and **Figure 2**Aii). Once all the landmarks are marked, we apply Imaris’ drift correction tool to align the scans. While this step effectively corrects most positional discrepancies, minor residual drift between scans may remain.

Following alignment, we proceed with lesion segmentation using the “Surfaces” tool (**Figure 1**Bv). For this, the operator determines a specific threshold to track the lesions in each scan to obtain the most accurate identification of lesion boundaries. Imaris then generates lesion objects based on this threshold and several other segmentation parameters (e.g., smoothing). Manual curation is required at this stage to remove false positives, such as vertebrae that often emit a PET signal and may be incorrectly labeled as TB lesions. Also, we found it useful to explore the scans using x/y/z slices both in VivoQuant and Imaris to make sure that all lesions identified in Imaris can also be located with VivoQuant. After accurately segmenting and manually correcting the lesions, the lesions could be grouped depending on their location in the thoracic cavity. For example, in many (but not all) scans we found heart to be highly labeled (e.g., **Figure 2**); in this case, heart-associated signal is stored in Imaris as “heart” Surface. Finally, characteristics of identified Surfaces (and landmarks) such as their channel intensity, volume, surface area, 3D coordinates, etc. are then exported into.xls (MS Excel) file for further analysis (**Figure 1**vi). In addition, Surface objects can be exported in a Virtual Reality file format (.wrl) that represents each lesion as a collection of points on the surface that can be also further analyzed (not shown in **Figure 1**B).

To rigorously compare characteristics of the lesions provided by Imaris (e.g., maximal intensity in the PET channel, *I*_max_) with the standard metrics generated by DICOM viewer VivoQuant (e.g., SUV_max_) we analyzed PET/CT data from four animals including two from a recent study^34^. In the published study^34^, animals were infected with a low dose of Mtb strain CDC1551 Mtb and then some of them were coinfected with SIV at 9 wks post Mtb infection (see Materials and methods for more detail). The animals had different pattern of developing active disease/tuberculosis with animal 41883 showing active disease starting with the first scan (wk 11) while animal 40884 did not progress to TB even after 18 wks.

We then followed our pipeline (**Figure 1**) to rigorously analyze the scans for the 4 animals (12 scans in total) both in VivoQuant and in Imaris. We paid close attention to label (Invicro) or automatically segment (Imaris) all lesions, so their basic characteristics (SUV_max_ and *I*_max_) could be rigorously compared. Importantly, the surfaces created using Imaris closely resemble the shape, size, and boundary lines of those generated by manual labeling in VivoQuant for all four animals regardless of whether the scan is rendered in 3D view (**Figure 2**A), or in sagittal or coronal slice views (**Figure 2**B&C and **Supplemental Figures S1 and S2**). This result supports the hypothesis that the findings from tracking PET/CT tuberculosis scans in NHP lungs using DICOM viewers can be effectively replicated using Imaris software.

It is interesting to note that heart is one organ that often produces very high intensity signal, well detectable in VivoQuant and Imaris, especially in 3D view (**Figure 2**). Because heart-derived PET signal is not thought to be important for TB pathogenesis, SUV_max_ for heart is typically excluded from the analyses. Interestingly, heart may not be visible in coronal view when a specific plane/slice is used to generate the image (**Figure 2**Biv-vi). Finally, even though the heart usually produces a signal, which was the case for most animals and scans analyzed, it is also possible for the heart to not produce a signal, as it occurred with animal 44104 at week 11 (**Supplemental Figure S2**Bi) and animal 41634 at week 18 (**Supplemental Figure S2**Ciii). The reason for this phenomenon remains unclear, although we speculate that it could be related to the animal’s diet or physical activity in the hours prior to the scan.

Imaris’ default color scale features a solid color. However, there is an option to switch to the “fire” color scale to closely match Invicro’s palette. For the images presented here, they are displayed using Imaris’ default solid color scale, specifically with a red selection in this instance. When comparing the rendering of the PET/CT scans in Invicro vs. Imaris, we observed that the intensity of the PET scans in VivoQuant is just as strong as that seen in Imairs. However, the PET signal in animal 40884 appears dimer in both VivoQuant and Imaris (compare **Supplemental Figure S1**Diii and **Supplemental Figure S2**Diii). The explanation for this remains unsettled, although we hypothesize that this could have been caused by a delay in the scan acquisition post-radiotracer injection, causing a larger portion of it to decay compared to other scans.

Another important observation we made when comparing scans analyzed in VivoQuant versus Imaris was the alignment of lesions and anatomical structures across multiple time points. For example, in animal 41883 there is a noticeable shift in anatomical structures between scans displayed in VivoQuant; specifically, when examining the heart’s position across different time points in the Invicro-processed scans, if one were to draw a horizontal reference line across the three images, the heart appears lower in the z-stack at weeks 11 and 16, and higher at week 22 (**Supplemental Figure S1**Ai-iii). However, when analyzing the same animal using the scans processed in Imaris, applying the same idea of the horizontal reference line reveals that the heart remains at a consistent position in the z-stack across all time points (**Supplemental Figure S2**Ai-iii). In contrast, when analyzing the VivoQuant versus the Imairs-processed scans for animal 40884, we see that for the three time points in the Invicro-processed scans, the alignment is actually fairly consistent and no significant shift is noted in the heart’s position across the z-stack (**Supplemental Figure S1**Di-iii). This indicates that while VivoQuant can achieve a satisfactory level of alignment between different time points for an individual animal without using landmarks, this method is less reliable and less effective compared to Imaris, which employs landmark positioning and drift correction. Indeed, by drawing this imaginary horizontal line we noted no apparent shift across the different time points for any of the animals as viewed in Imaris (**Supplemental Figure S2**). This consistency highlights Imaris’ key advantage of its ability to correct for drift and align scans from different time points. This alignment allows for more accurate comparisons and the tracking of lesion/sites of inflammation over time, ultimately enabling for more rigorous characterization of disease progression.

### Maximum channel intensity in Imaris matches well SUV_max_ in VivoQuant

One of the main challenges in translatability of the analysis of PET/CT scans from DICOM viewers to Imaris was the lack of a feature or algorithm in Imaris capable of automatically converting channel intensity units to SUV, which is the standard metric used to measure FDG radiotracer absorption in various lung lesions. The difficulty is in units – when DICOM files are imported into Imaris, the PET intensity is converted to channel intensity with the maximum 32,767 (16 bit color) and no information about the injected dose or animal weight is imported into Imaris.

To account for the FDG dose injected and the weight of the animal, for each lesion we generated a normalized maximum channel intensity *nI*_max_ that scales the actual maximal intensity *I*_max_ by the injected dose, weight of the animal, and a scaling factor *f* (**eqn. (1)** and see Materials and Methods for further details). The number of lesions varied between the animals, with animal 41634 having only *n* = 10 lesions while animal 44104 having *n* = 53 lesions (**Supplemental Figure 3**). Of note, we considered the lesions identified in different scans as independent even though it is likely that some of the lesions are the same at different time points (**Figure 2**). Interestingly, we found excellent correlation between normalized maximum channel intensity *nI*_max_ and SUV_max_ for each of the lesions at any given scan/time point; these are characterized by the slope *m* between the two metrics with *m* = 1 corresponding to perfect correspondence (**Figure 3**).

**Figure 3:**
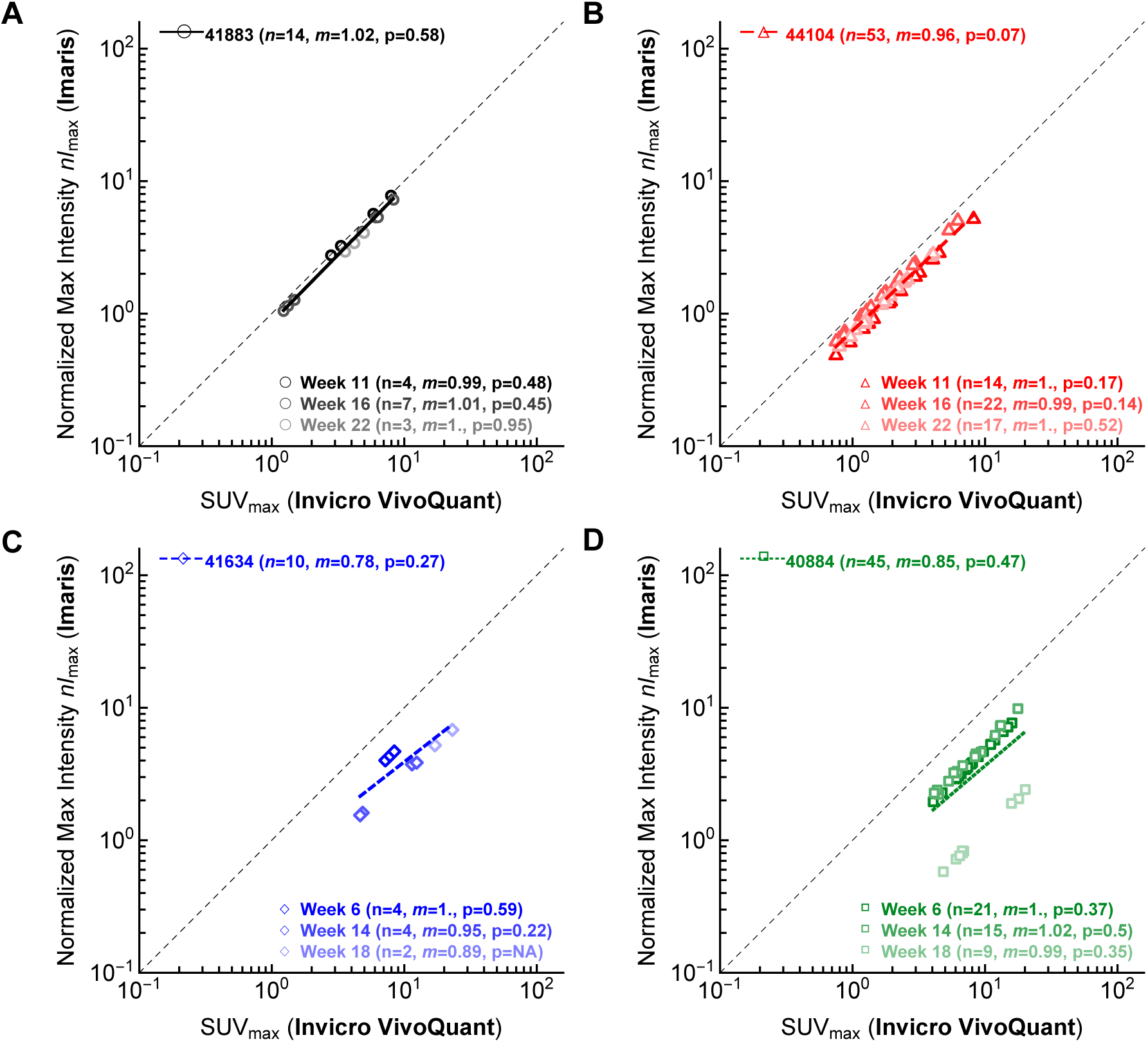
Normalized maximum PET channel intensity, determined by Imaris, matches well standard max SUV provided by VivoQuant. We analyzed PET/CT data from four Mtb-infected RMs (each with three PET/CT scans) either with VivoQuant, a standard software for PET/CT data analysis, or Imaris (Bitplane), software typically used for analysis of fluorescent microscopy data. For each lesion we plot maximum SUV as provided by VivoQuant (x axes) or maximum intensity as provided by Imaris (y axes), normalized using weight of the animal and 18F-FDG injection dose and multiplied by *f* = 10^4^ (see **eqn. (1)**). We show data for animal 41883 (**A**), 44104 (**B**), 41634 (**C**), and 40884 (**D**) with *n* indicating the number of lesions detected in all scans and *m* being a regression slope between SUV values of VivoQuant vs. Imaris for all lesions or lesions found in individual scans; we show the regression lines in individual panels for all lesions. Scans done at different times are shown by different color intensity. The dashed line shows the slope of 1.

While the measurements appear to cluster along a single line in animals 41883 and 44104, there is a clear discrepancy between *nI*_max_, detected in Imaris, and SUV_max_, detected in VivoQuant for scans taken at different times (**Figure 3**C&D). For animal 41634 the *nI*_max_ for week 6 was higher than at weeks 14 and 18, while for animal 40884 the *nI*_max_ for week 18 was lower than at weeks 6 and 14. This phenomenon is further evidenced in raw scans revealing that the week 18 scan for animal 40884 appears dimmer than those of earlier time points in both VivoQuant and Imaris, respectively (**Supplemental Figure S1**Di-iii and **Supplemental Figure S2**Di-iii). Given that this decrease in perceived intensity was observed in both VivoQuant and Imaris-processed scans, yet the SUV_max_ obtained from VivoQuant did not exhibit the same deviation noted in the calculated *nI*_max_ from Imaris, it suggests that VivoQuant might utilize an internal algorithm to compensate for this visually perceptible reduction in intensity when calculating SUV, thus preventing discrepancies in the resulting SUV values. This may highlight a limitation of our methodology and emphasize the importance of interpreting the results with care. It also underscores the need for additional studies to refine this methodology, making it more broadly applicable in future research. Furthermore, the scan of week 6 of animal 41634, did not visually show such a remarkable change in perceived intensity, which aligns with the smaller deviation in *nI*_max_ compared to the more pronounced variability seen in animal 40884. Consequently, a noticeable change in intensity would be difficult to detect in this case.

Although the normalization of the maximal intensity of PET signal generated by Imaris for individual lesions by animal weight and injected radiotracer dose seems to be important, this is an extra step one needs to do – i.e., to extract the dose/weight information from metadata of DICOM files. We therefore investigated if simply using actual maximum PET channel intensity, provided by Imaris for individual lesions, may nevertheless provide important information. Interestingly, we found excellent correlation between the actual maximum intensity of PET signal for individual lesions *I*_max_, as provided by Imaris, and SUV_max_, generated by VivoQuant (**Supplemental Figure S3**) for individual scans and over time. While the absolute values of maximal intensity are high, they scaled linearly with SUV_max_ suggesting utility of this approach. Yet, more analyses will need to be done to determine if indeed simply using maximal intensity in PET channel would be sufficient to indicate cellular activity in a lesion, especially given that the body weight does impact estimated SUV_max_ ^51^. It is typical, though, to adjust the injected FDG dose by the weight of the animal and the time between radiotracer production and imaging, thus, potentially removing variability in PET signal that comes from the body weight.

### Using Imaris-derived characteristics to track disease development

One important fea-ture of Imaris is various characteristics it generates for each of its objects such as their location, volume, surface area, etc., and relationship between objects (e.g., distance to the closest another object). By tracking these characteristics over time we can build better understanding of how the animals progress towards active disease and/or control the infection (**Figure 4**). In our cohort of four animals, animals 44104 and 41634 did not exhibit active disease at the first scan post-infection but developed active disease at later time points. In contrast, animal 41883 had active disease at all scans and animal 40884 had only evidence of Mtb infection (formerly denoted as latent TB infection^52^) and no active disease.

**Figure 4:**
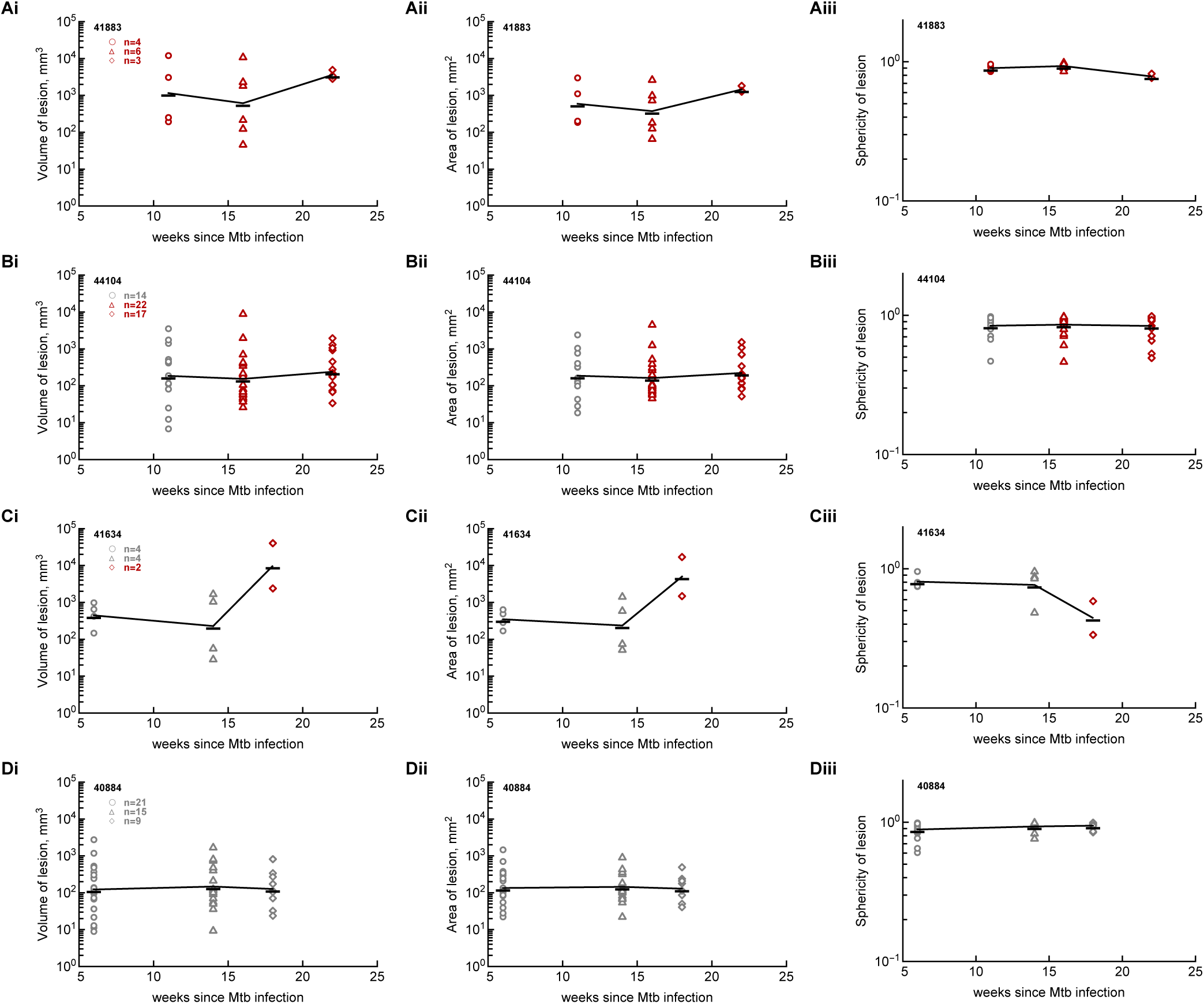
Tracking volume, area, and sphericity of PET lesions identified by Imaris. For every lesion identified for animals 41883 (**A**), 44104 (**B**), 41634 (**C**), and 40884 (**D**) we show lesion’s total volume (**i**), surface area (**ii**), and sphericity (**iii**). Markers denote individual lesions and lines connect average values per time point; color of the markers denotes whether the animal was classified as being asymptomatic (**gray**) or having TB (**dark red**) at specific scan times.

Interestingly, we found that the mean volume for the lesions of each animal remained fairly constant, with little to no variability across the different time points and within the range of approximately 100–2,000 mm^3^, **Figure 4**i); a notable exception is the final time point for animal 41634, which exhibited a notably larger mean lesion volume (**Figure 4**Ci). Given actual PET scans, we hypothesize that large rise in the volume of the lesions in this animal is due to their merger leading to inflammation of the whole left lung (**Supplemental Figure S1**C and **Supplemental Figure S2**C). This could explain both the increase in lesion volume and the decrease in lesion count (**Figure 4**Ci). Furthermore, when examining the relationship between lesion volume and disease state, we observed that animal 41634 transitioned from an asymptomatic to a symptomatic state between weeks 14 and 18 (**Figure 4**Ci). During this period, there was a substantial increase in the mean volume of its lesions, increase in the surface area, and decrease in sphericity of the lesions (**Figure 4**C). In contrast, although animal 44104 experienced an earlier transition from a latent to an active disease state between weeks 11 and 16, there was no significant change in the mean volume, mean surface area, or mean sphericity of the lesions during that interval (**Figure 4**B). However, while the mean volume remained relatively constant, most lesions increased in size, effectively shifting the overall volume range upward. This observation highlights the importance of not relying solely on mean lesion volume as predictor of TB progression. Analyzing the distribution and range of lesion volumes may provide additional insight, particularly if a single lesion, especially one located in a critical anatomical region, undergoes substantial growth and serves as the primary driver of disease progression and symptom development.

We observed a similar pattern in the surface area values per lesion (**Figure 4**ii). Means for lesion areas remained constant, with little to no variability between the different time points and within a range of approximately 100–2,000 mm^2^, with the exception of the final time point for animal 41634. Finally, we also examined the same phenomenon when correlating surface area with changes between disease states. We also noticed when animal 41634 transitioned from an asymptomatic to a symptomatic state between weeks 14 and 18 (**Figure 4**Cii), there was also a noticeable increase in the mean area of its lesions, suggesting a possible correlation between mean surface area and the transition to an active disease state that would have to be further investigated. We also noted, as with volume per lesion, that although there was no significant change in the mean surface area when animal 44104 transitioned from a latent to an active disease state (**Figure 4**Bii), most lesions did increase in size, effectively shifting the overall volume range upward. This observation further underscores the importance of not solely focusing on the mean, but that analyzing the distribution and ranges may provide additional insights into the dynamics of disease progression.

Lastly, we analyzed changes in lesion sphericity as a measure of lesion shape. Most lesions had mean sphericity of 0.7 or higher, indicating generally spherical shapes (sphericity of 1 indicates a perfect sphere). The exception was again found in the final time point for animal 41634, where the mean sphericity dropped to approximately 0.45. This lower value indicates that the lesions were less spherical, further supporting the hypothesis of inflammation of the whole left lung of this animal. This finding supports the idea that morphological characteristics such as sphericity may serve as an indicator of disease progression. Moreover, from the decline in sphericity in animal 41634, we can hypothesize that changes sphericity values may reflect structural lesion changes associated with pathology (whole left lung inflammation in this case).

Another important observation was that animals with stable lesion size and shape across multiple time points remained asymptomatic, such as animal 40884 (**Figure 4**Di-iii). The means for volume, surface area and sphericity remained fairly constant over time for this animal, and the distributions for each of these metrics also showed minimal variability. In contrast, animals that transitioned from a latent to an active disease state displayed more dynamic changes. For example, in animal 41634, there was a noticeable increase in both mean lesion volume and surface area, accompanied by a decrease in sphericity, suggesting a shift toward more irregularly shaped lesions (**Figure 4**Ci-iii). Similarly, in animal 44104, although the mean values for volume and surface area did not change significantly, the ranges of their distributions shifted upward, indicating that most lesions increased in size. However, this animal did not exhibit changes in sphericity (**Figure 2**Bi-iii). This finding suggests that these three metrics could provide valuable insights into the TB progression and play a significant role in understanding the transition between different disease states.

Even though both VivoQuant and Imaris allow to visualize location and PET intensity of individual lesions (e.g., **Figure 2A**), we developed methodology to graphically display 3Dnlocation, size, and PET intensity of the lesions using information provided by Imaris for individual Surfaces. Specifically, we converted the volume of each lesion *V* into spheres with the radius 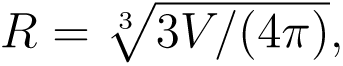, and positioned these “spheres” using lesion’s 3D coordinates provided by Imaris. In addition, we used hue levels to indicate scaled normalized maximal PET intensity (*nI*_max_) in each lesion, and used colors (black vs. red) to indicate disease status of the animal (**Figure 5** and **Supplemental Figure S4**). This representation of lesions in each animal thus allow to visualize dynamics of lesions over time in 3D.

**Figure 5:**
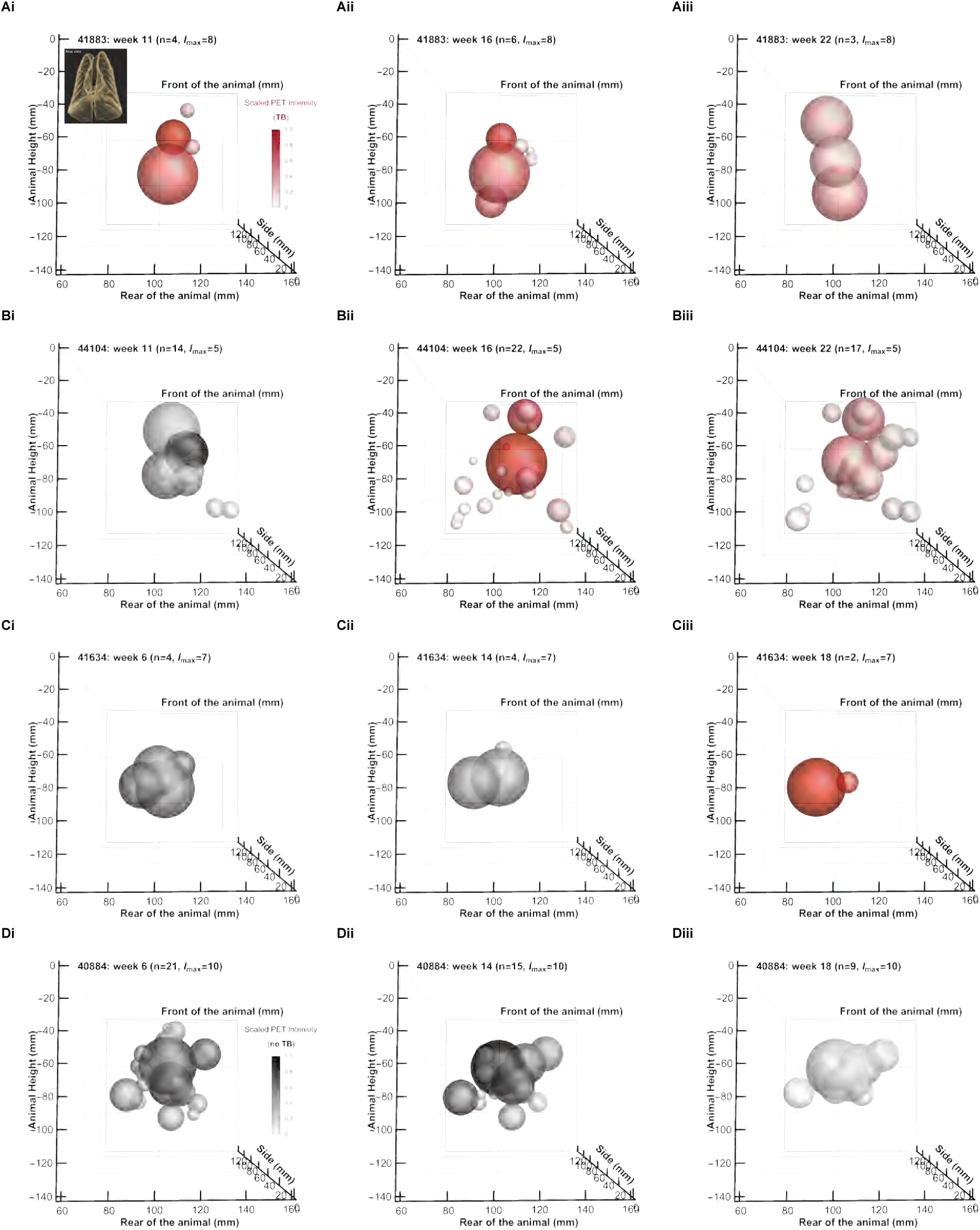
3D rendering of individual PET-defined lesions in Mtb-infected monkeys using metrics provided by Imaris: rear view. By using XYZ coordinates, provided by Imaris we plotted individual lesions detected in animals 41883 (**A**), 44104 (**B**), 41634 (**C**), and 40884 (**D**) at different time points after infection (times are denoted on individual panels). We show coronal view of the animals with rear of the lung being closer to the viewer (see also **Supplemental Figure S4**). On z-axis we show the height of the animal starting at the apex of the lungs. We show the total number of lesions detected at each time point as *n*. Lesions are plotted as spheres with the radius 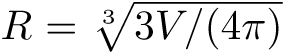 where *V* is the PET volume determined by Imaris (**Figure 4**). Intensity of the color denotes scaled maximum PET signal intensity normalized to *I*_max_ values for the largest maximum intensity for all lesions in all scans of a given animal; this also takes into account the actual injected FDG dose and weight of the animal (**Figure 3**). In panel Ai the insert shows the coronal (rear) view of the lung segmentation of a representative animal. Black colors indicate animals that were asymptomatic and red colors indicate that the animal was diagnosed with active disease (TB).

While visual representation of lesions is exciting, 3D rendering comes at a cost of perceived size of lesions as lesions that occur closer to the viewer will appear larger than perhaps larger lesions that are located farther. For example, the largest lesion at week 18 for animal 41634 (**Figure 5**Ciii) may appear similar in size to the two largest lesions from week 14 for the same animal (**Figure 5**Cii). However, the radii of the two largest lesions from week 14 are 6.3 mm and 7.4 mm, whereas the radius of the largest lesion at week 18 is 21.3 mm. Also, the center of mass for the 18-week lesion is positioned further back, affecting its visual representation due to our perception of depth.

When examining the number and volume of lesions, we observe an inverse correlation between lesion count and volume. Specifically, a decrease in the number of lesions between two time points is generally accompanied by an increase in most lesions’ volume (**Figure 5**). For example, in animal 41883, the number of lesions (*n*) decreases from 6 to 3 between weeks 16 and 22, respectively; correspondingly, the lesions at the later time point appear larger (**Figure 5** and **Figure 4**). This trend is evident not only in symptomatic animals such as 41883 but also in asymptomatic ones like animal 40884, where a reduction in lesion count is observed across all three time points and is also accompanied by seemingly larger volumes of the remaining lesions (**Figure 5** and **Figure 4B**).

We noted an inverse pattern in animal 44104, which demonstrates an increase in lesion count from 14 to 22 between weeks 11 and 16, despite having smaller overall lesion volumes (**Figure 5**Bi-ii). This rise in lesion count coincided with a transition from asymptomatic state to active disease, suggesting a possible correlation between increased lesion count and decreased lesion volume upon symptom onset. However, this interpretation is challenged by observations in animal 41634, where the shift to an active disease state between weeks 14 and 18 was accompanied by a decrease in lesion count alongside a marked increase in total lesion volume (**Figure 5**Cii-iii), contrasting sharply with the trend seen in animal 44104. Such differences in granuloma dynamics are consistent with the well-documented heterogeneity of Mtb growth/elimination in individual granulomas of macaques^43,53^. It is conceivable that in animal 41634, Mtb is eliminated in some granulomas leading to resolution, while Mtb continues to grow in others leading to expanded granulomas, resulting in fewer but larger lesions. Together, these contrasting patterns highlight the complexity of granuloma behavior in NHPs and underscore the need to further elucidate the relationship between lesion dynamics and clinical disease progression.

By examining the darkness of the lesions at various time points, we observed that in the animals that did not show changes between disease states (41883 and 40884), the lesions gradually become dimmer over time. This may indicate a reduction in bacterial burden, suggesting a potential recovery in animal 41883 or a containment of the disease in animal 40884. In contrast, the lesions observed at week 18 in animal 41634 appear darker than those at earlier time points, reinforcing the likelihood that this animal developed inflammation of the whole left lung (**Figure 5**C). Similarly, the lesions in animal 44104 at week 16 also show increased darkness compared to week 11 (**Figure 5**). The emergence of darker lesions in animals 41634 and 44104 at the first time points when they exhibited symptoms raises the possibility that an increase in *nI*_max_ may be correlated with the onset of symptoms and, therefore, with the activation of tuberculosis. Lastly, it is notable that in most scans, the lesions with the highest *nI*_max_ values are located centrally. Given that these images present the rear (posterior) view of the animals, and by analyzing this figure in conjunction with front view images (**Supplemental Figure S4**), it appears that these centrally located lesions are likely situated in the mediastinum. This suggests that the lesions exhibiting the highest maximum intensities correspond to thoracic/lung-draining (mediastinal) lymph nodes^54^. We believe this methodology could be valuable for investigating the correlation between *I*_max_ and/or SUV_max_ with the lesions’ anatomical location and thus may hold potential as a predictive tool for assessing disease progression.

## Discussion

DICOM viewers such as VivoQuant or OsiriX are widely used for the general visualization and analysis of PET/CT scans in TB studies involving NHPs ^34,39^. However, analysis of PET/CT scans, for example, with VivoQuant involves a tedious and time-consuming pipeline that requires extensive scrolling through the different 2D slices across the different anatomical planes (sagittal, coronal and transverse) and the manual segmentation of the sites of inflammation (**Figure 1**). By developing a novel pipeline we found that Imaris provides numerous benefits in this regard, as it streamlines the workflow, reduces the time spent in image analysis and provides a more rigorous technique for lesion segmentation (**Figure 1**). In our experience, segmenting the lesions based on an intensity threshold in Imaris rather than manually contouring the ROIs in VivoQuant shortens the time spent on the analysis of a scan by about 66% (1 scan in VivoQuant vs. 3 scans in Imaris per day), with further speed improvements achieved when serial scans of the same animal are analyzed in one setting. This results in an increase of the output of scans analyzed. Additionally, although the intensity threshold would still have to be determined by the operator, this value, along with the other parameters set in the creation wizard of Surfaces in Imaris, can be standardized and recorded, making tracking and documentation a more rigorous process. This enhances reproducibility and facilitates sharing and replication of results in future studies. Moreover, Imaris provides a comparably larger amount of quantitative metrics compared to VivoQuant. While VivoQuant typically provides basic values for a lesion such as maximum, minimum, and total SUV, volume, and HUs, Imaris also exports morphological descriptors (e.g., sphericity, ellipticity), spatial coordinates (e.g., center of mass and boundary points), and channel intensity data from both PET and CT channels. These additional metrics offer new opportunities to explore lesion characteristics and their potential correlation with TB progression, as well as enhance our ability to better understand TB heterogeneity at the level of the granuloma.

We demonstrated that we can effectively track the same lesions identifiable in VivoQuant with Imaris (**Figure 2** and **Supplemental Figures S1 and S2**). Additionally, Imaris offers the critical advantage of aligning scans across multiple time points using anatomical landmarks. This functionality has significant potential for tracking lesion evolution, including merging, splitting, or slight changes in location over time. The ability to confirm lesion continuity based on spatial location, morphology, and intensity values could yield deeper insights into TB progression dynamics.

We also found that the normalized PET channel intensity values obtained using Imaris correlates well with the standardized metric of radiotracer uptake/metabolic activity used in the field such as SUV_max_ (**Figure 3**). Interestingly, even non-normalized, raw PET channel intensity as provide by Imaris, correlated extremely well with SUV_max_ (**Supplemental Figure S3**); we believe that this is because in our scans, FDG injected dose is typically adjusted to the weight of the animal and the timing of the scan relative to ^18^F-FDG production. Additional Imaris-derived metrics such as volume, surface area, and sphericity may also offer valuable insights into TB pathology, particularly in relation to transitions between latent and active disease states (**Figure 4**). Anatomical location of the lesions and their correlation with *nI*_max_ (e.g., **Figure 5** and **Supplemental Figure S4**) could help future studies investigating not only TB progression but also drug delivery mechanisms in granulomas.

Due clinical applicability of PET/CT scanning technologies, there have been much research to standardize generation and analysis of PET/CT data ^55–57^. There have been many methods to improve representation of PET signals to help better diagnose patients^57^. There also have been novel advances, including with using machine learning, to rigorously quantify PET images, especially in cancer research^58,59^. In TB research, however, manual labeling of PET-defined lung lesions remains relatively common. Using more advanced segmentation techniques, as those available in Imaris, could help to more rigorously determine how location and types of lung inflammation, caused by Mtb, results in active disease.

Our work has several limitations. Imaris is a proprietary and expensive software that may not be accessible in all research institutions; in contrast, DICOM viewers while also being expensive, are often bundled with PET/CT scanners. While Imaris streamlines segmentation using threshold-based automation, some degree of operator bias remains, as thresholds must be manually defined, and some identified Surfaces need to be manually removed (e.g., spine). Additionally, even though lesion’s normalized maximum PET channel intensity *nI*_max_, provided by Imaris, correlates well with SUV_max_ for the same lesion, we acknowledge that *nI*_max_ or raw value of the channel intensity *I*_max_ is not a direct replacement of SUV_max_. Converting raw intensities *I*_max_ to normalized intensities *nI*_max_ requires a certain degree of computational expertise, adding complexity to the workflow. Similarly, the raw intensity of the CT channel also needs to be proceed to convert them into commonly accepted Hounsfield units (**HU**). The meta data information of DICOM files such as intercept and slope should be helpful in this regard.

Another consideration is the anatomical landmark selection used for drift correction and scan alignment in this methodology. While these landmarks were chosen in consultation with veterinary experts to ensure their stability and consistency over time, this is the first application of such a method, and its generalizability and replicability remain to be tested. In particular, when working with additional PET/CT scans we found that sometimes vertebra T2 or T10 are missing from the scans, due to the positioning of the animal on the scanner’s bench. Whether using vertebrate T3 and T9 for location of landmarks would be more advantageous remains to be determined. We found it perplexing that while there may be overall reduction in total PET intensity detected in one scan (and shown in Imaris) as compared to scans done at other times (**Supplemental Figure S1**D and **Supplemental Figure S2**D), VivoQuant can effectively compensate for it when calculating SUV_max_ but Imaris does not (**Figure 3**D). The full relationship between the injected dose, animal weight, overall metabolic state and the total PET intensity in the scan will need to be determined. Finally, although Imaris can export 3D boundary coordinates of Surfaces as virtual reality files (.wrl), a standardized method to track lesion identity across time points and to correlate features of these Surfaces to disease progression remains to be developed.

Our study opens avenue for future research. First, we aim to develop a robust approach for tracking lesion identity over time using Imaris-exported virtual reality files (.wrl) and associated morphological and spatial metrics. This would improve our ability to characterize lesion evolution and its correlation with transitions between active and latent disease states. Second, adapting our pipeline to convert CT intensities into HU and finding robust way to convert PET channel intensity to SUV would make Imaris more functionally comparable to DICOM viewers; using metadata in DICOM files should be helpful in this process. Third, we aim to explore the feasibility of using Imaris’ machine learning tools to segment TB lesions using either PET/CT data or CT data alone; the latter could reduce dependence on PET imaging—a modality that is less widely available and more costly than X-ray or CT imaging. We are also exploring if using more landmarks to more accurately align animal scans may improve precision to accurately track development of individual lesions over time. Using automated Sufrace-tracking tools in Imaris may be particularly useful. Lastly, expanding this methodology to open-source or lower-cost softwares with similar capabilities could greatly increase its accessibility and impact in the field. Doing so would allow more laboratories to perform high-throughput, rigorous TB lesion analysis while generating deeper insights into disease progression and drug delivery or therapeutic response. Finally, increasing the size of the animals and scans should help to better define features of the PET-derived lesions that could explain why some animals progress to active disease while others are able to control (and perhaps clear) the infection.

## Data sources

The data from the paper (digitized from original publications) along with the codes are available on github: https://github.com/vganusov/pet-ct-imaris-vs-invicro

## Code sources

All analyses have been primarily performed in Mathematica (ver 12.1)

## Ethics statement

No new animal work has been performed.

## Author contributions

VVG conceived the overall concept of the study. DK and SM provided experimental data from their studies and guided interpretation of experimental observations. XA helped with training EH in imaging data analysis in Invicro VivoQuant. EH performed analysis of PET/CT scans in Invicro VivoQuant and Imaris, and developed the pipeline of PET/CT data processing in Imaris. EH wrote the first draft of the paper and all authors read, edited, and agreed on the final version.

## Acknowledgments

We would like to acknowledge the Imaris customer support team for their guidance in exploring the analysis of PET/CT scans using this software package; Dr. Corinna Ross and Ms. Kathi Darrow for their assistance in identifying anatomical landmarks in NHPs to align the scans, and Manuel Aguilar for outlining the complex procedures and nuances involved in performing PET/CT scans in NHPs at Texas Biomed. We also thank Pauline Maiello, Dr. Joanne Flynn, and all other members of their research group for sharing data that, although not included in this study and yet to be analyzed, provided valuable insights into the structure and interpretation of PET/CT scans and DICOM files.

## Financial Disclosure statement

This work was supported in part by the NIH/NIAID grant R01AI158963 to VVG, grants R01AI111914, R01AI134240, R01AI138587, R01AI135729, R01AI135751, R01AI150043, and R01AI155346 to DK and grants R01AI134245 and R01AI181701 to SM. In addition, this research has been facilitated by the infrastructure and resources provided by the Texas Biomedical Research Institute Interdisciplinary NexGen TB Research Advancement Center, an NIH funded program (P30AI168439).

## Abbreviations

PET: positron emission tomography
CT: computed tomography
TB: tuberculosis
Mtb: *Mycobacterium tuberculosis*
NHPs: nonhuman primates
LNs: lymph nodes
SUV: standardized uptake value
DICOM: Digital Imaging and Communications in Medicine
FDG: Fluorodeoxyglucose
SIV: simian immunodeficiency virus
ART: antiretro-viral therapy.

## Supplemental Information

**Supplemental Figure S1:**
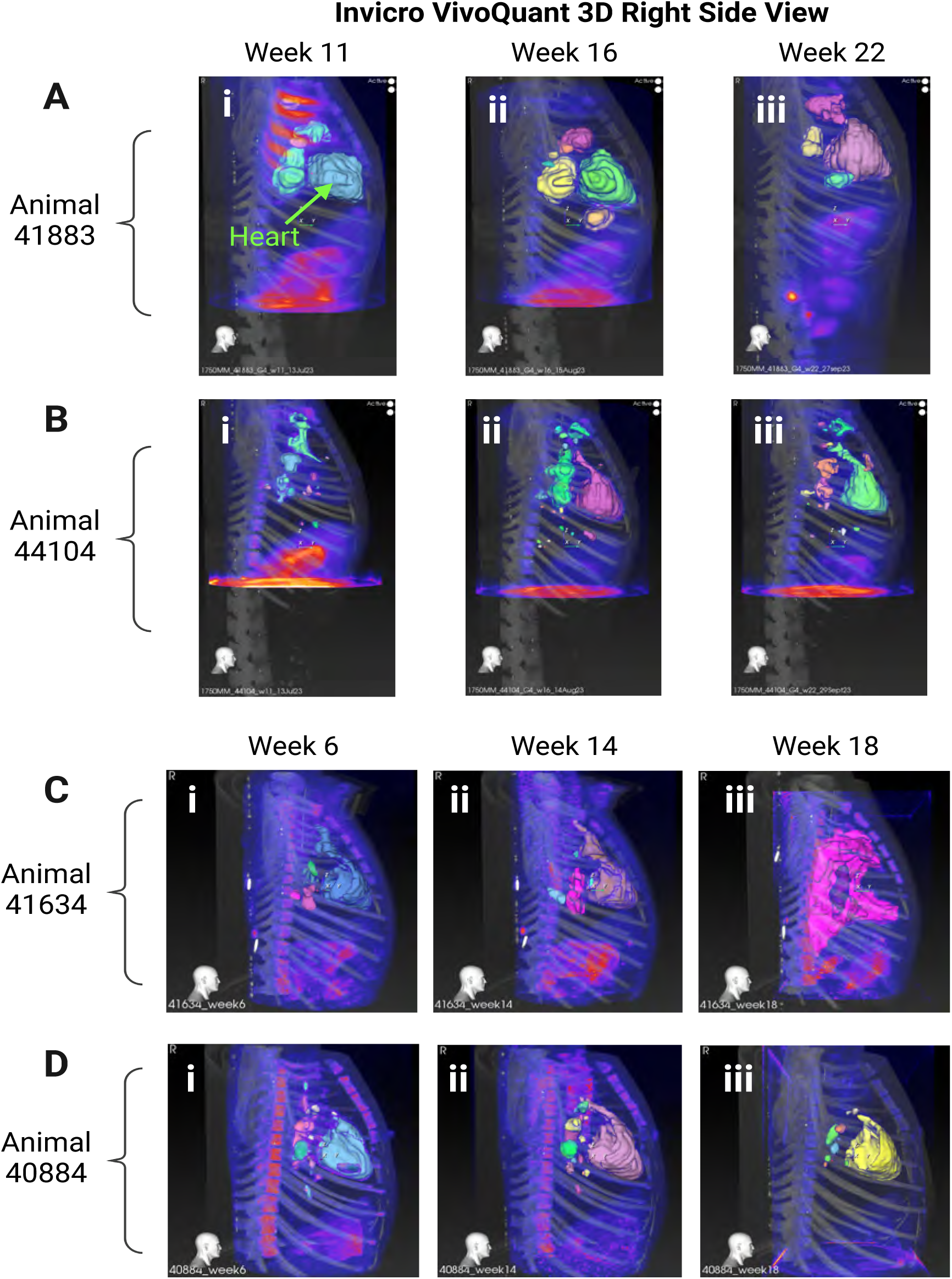
**All scans for all animals and all time points processed in Invicro VivoQuant**. We show lesions identified for all animals using VivoQuant at different times post-infection from the right side 3D view for animals 44883 (**A**) and 44104 (**B**) at at 11 **(i)**, 16 **(ii)** and 22 **(iii)** weeks post-infection; and for animals 41634 (**C**) and 40884 (**D**) at 6 **(i)**, 14 **(ii)** and 18 **(iii)** weeks post-infection. PET signal associated with the heart (shown by an arrow in **Ai**) was excluded from our analyses.

**Supplemental Figure S2:**
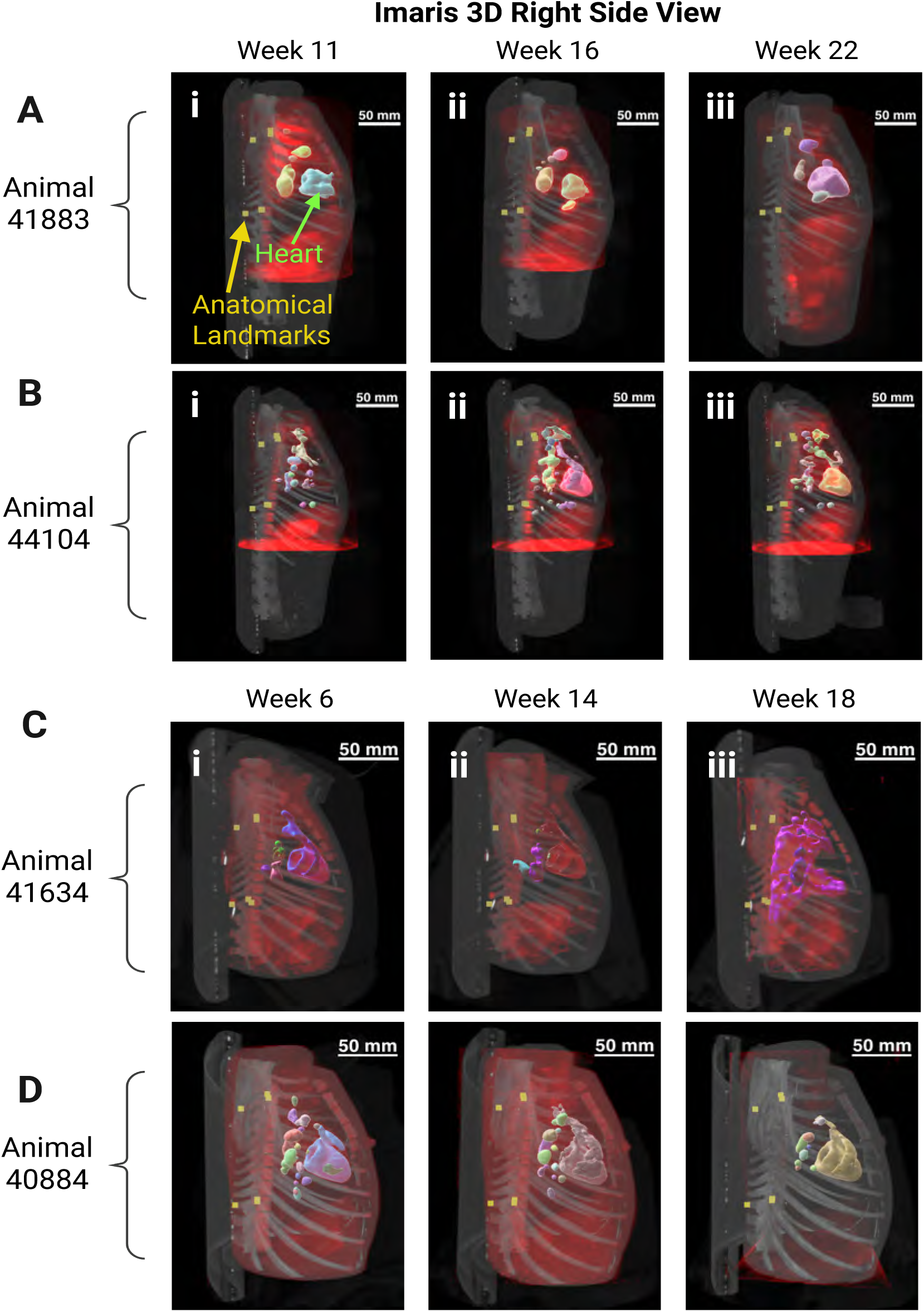
**All scans for all animals and all time points processed in Imaris**. We identified and quantified the same lesions as in **Supplemental Figure S1** with Imaris.

**Supplemental Figure S3:**
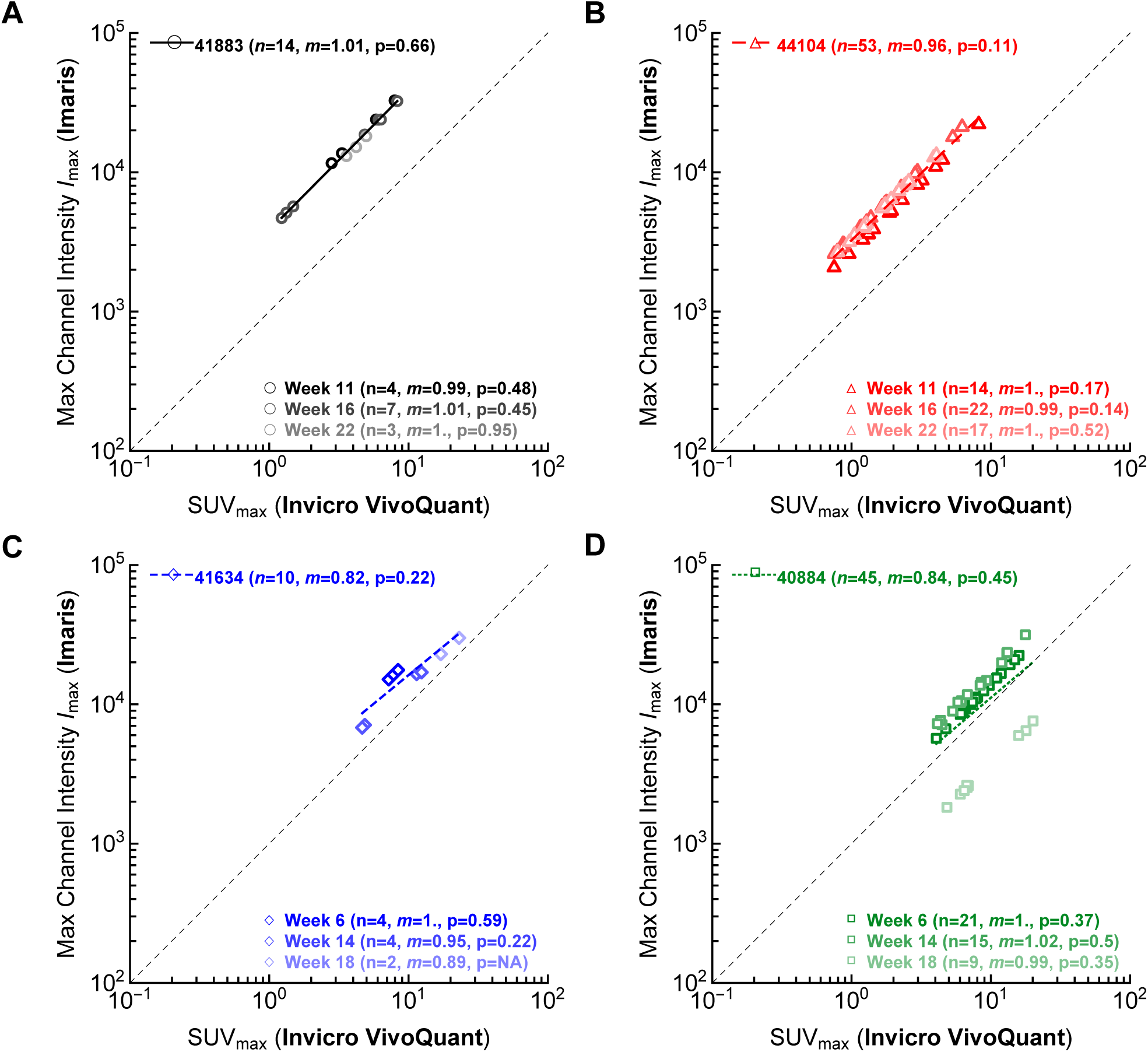
**Maximum PET channel intensity, determined by Imaris, also matches well max SUV provided by VivoQuant**. We performed similar analysis as in **Figure 3** except we used the raw values of maximum intensity *I*_max_ provided by Imaris (i.e., we ignored scaling given in **eqn. (1)**). Other analyses and notations are similar to those in Figure 3.

**Supplemental Figure S4:**
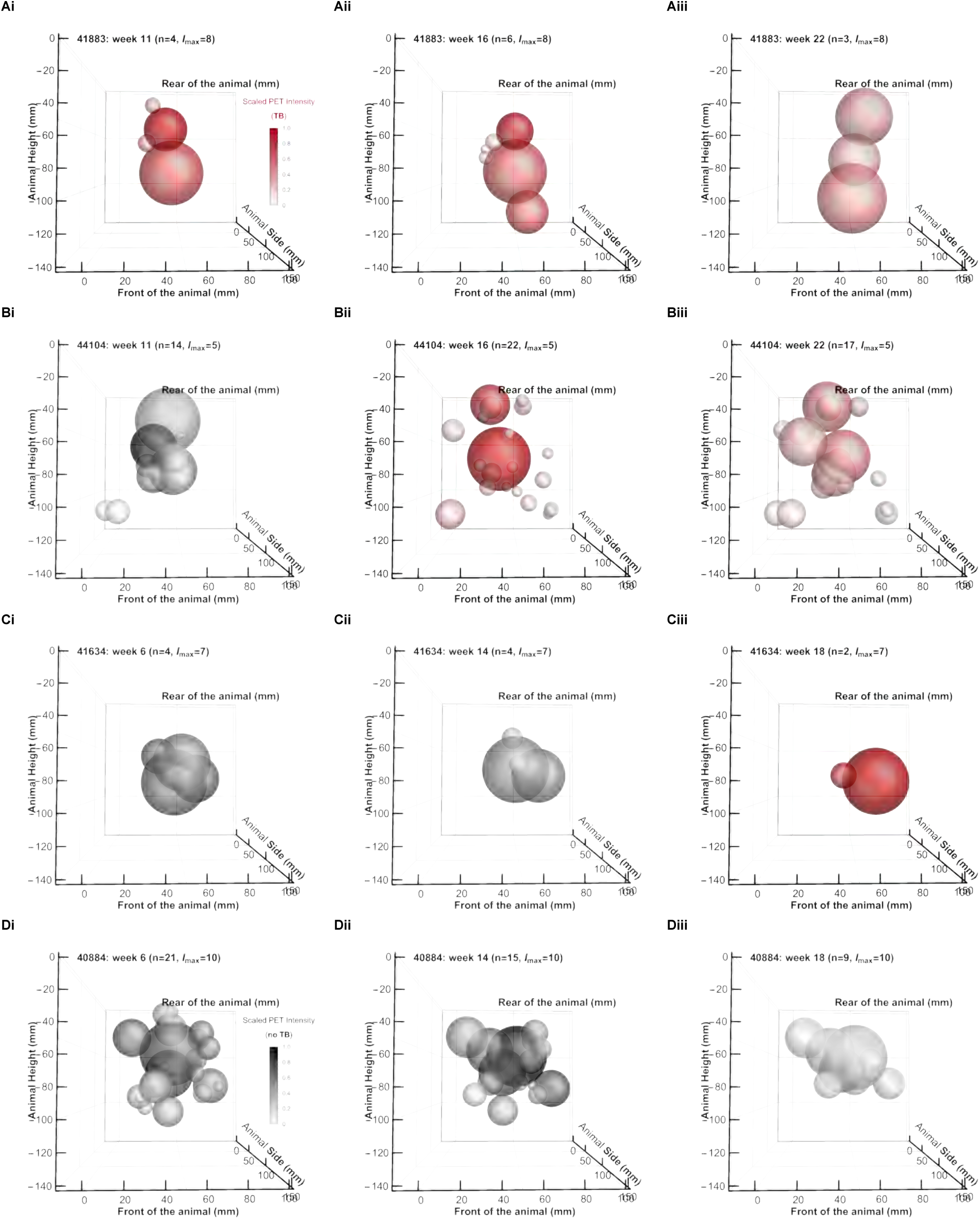
**3D rendering of individual PET-defined lesions in Mtb-infected monkeys using metrics provided by Imaris: front view**. The same data are plotted as in **Figure 5** for coronal view of the animals with front of the animal/lung facing the viewer.

## Notes

### Competing Interest Statement

The authors have declared no competing interest.

### Summary of Updates

We updated the materials and methods to improve clarity (per reviewers' suggestions); we slightly edited some figures to improve their redability. Add some additional points for Discussion.

